# A Short 63-Nucleotide Element Promotes Efficient circRNA Translation

**DOI:** 10.1101/2025.05.13.653105

**Authors:** Martina Chiara Biagi, Andrea Giuliani, Alessia Grandioso, Gaia Di Timoteo, Irene Bozzoni

**Author notes:** The authors wish it to be known that, in their opinion, the last two authors should be regarded as joint last authors.

## Abstract

Circular RNAs (circRNAs) are a class of RNA with multiple functions, including the ability to be translated.

Several intrinsic features of circRNAs, such as high stability, confer them advantages over linear RNAs; therefore, circRNA-based drugs have recently received increasing attention.

However, the inefficiency of their cap-independent translation and the difficulties in the large-scale production of long circRNAs negatively impact on their use in therapy. Some efforts have been done to solve these issues related to circRNA adoption, but, to date, circRNA translation still relies on long IRESs (600-800) and chemical group addition. In this study, identified a 63-nt element able to drive circRNA translation comparably to the most commonly used IRESs. This element includes a a 13-nt sequence previously reported to enhance linear RNA translation and a segment of the UTR of the endogenously translated circRNA circZNF609. Notably, this element combines a comparable IRES-like efficiency to a considerably shorter length, expanding the landscape of ORFs potentially suitable for being translated from circRNAs and enhancing their potential as therapeutic agents in therapy.

## Introduction

The resourceful nature of RNA has made it very appealing for therapeutic purposes. In recent years, different types of RNA have been employed in therapy such as RNA aptamers, antisense RNAs, small interfering RNAs or guide RNAs^1^.

mRNA based drugs were the first to be tested, given their direct activity on protein production: using an mRNA molecule carrying the correct sequence to rescue a mutation or to increase protein production is the most immediate solution to a genetic disorder or to trigger immune response in order to achieve vaccination.

Covid-19 made headlines for great diffusion and aggressivity, aside to the virulency it showed. Application of RNA received deserved attention during the pandemic due to the production of RNA-based vaccines.

However, the employment of linear RNA has its specific downsides: linear RNAs short half-life remains the main problem of RNA-based drugs.

Indeed, linear RNA is easily and rapidly degraded by exonucleases, which are present in the environment the RNA is released into (i.e. blood or serum)^2^.

Circular RNAs (circRNAs) have emerged as a fascinating class of non-coding RNAs, with their unique closed-loop structure conferring remarkable stability and resistance to exonucleases with respect to linear RNAs^3^.

While circRNAs were initially considered as mere byproducts of splicing errors, advances in high-throughput sequencing and computational biology have unveiled their pivotal roles in regulating gene expression and functioning as crucial players in various cellular processes^4^.

One of the most intriguing aspects of circRNA biology is their potential to encode functional peptides through a process that has been the subject of growing interest in recent years^5^. Unlike canonical linear mRNA, having no cap or poly-A tail, coding circRNAs utilize Internal Ribosome Entry Sites (IRES) to initiate cap-independent translation^6^.

IRESs are not the only structures we can employ in cap-independent translation. For example, circZNF609, one of the few examples of translated circRNAs, is translated from two alternative START codons also thanks to specific m^6^A modifications present in its untranslated region^7^.

The exploration of IRES elements within circRNAs has opened a new frontier in molecular biology, with implications spanning from fundamental cellular processes to potential therapeutic applications. However, our understanding of the diversity and functionality of IRES elements within circular context remains limited and limiting^8^.

Recently other 5’ UTR elements have been brought to general attention: Translation enhancing elements (TEEs). TEEs are 5’ UTR sequences rich in AUG triplets, positioned in cis upstream of a putative ORF able to boost protein production^9^.

Sequence analysis revealed the presence of a 13-nucleotide long motif that if added or removed from TEEs sequences could in fact enhance or reduce protein translation^10^. To date, different TEEs have been tested *in vitro* in a linear context^10^.

In this study, we efforted the exploration not only of diverse viral or human IRESs but also of other translation elements in order to test their translation regulation potential within the context of a circular GFP (cGFP) as well as of an endogenously translated circRNA, circZNF609^5^. We identified a small (about 60nt) but efficient element able to carry out circRNA translation and we engineered its sequence to increase its performance. Considering the growing popularity of single-stranded therapy, this is a significant achievement when you consider that the smaller the RNA to be circularized, the greater the circularization efficiency with current *in vitro* circularization methods. Eventually, we achieved a substantial minimization of the regulatory elements necessary for circRNA translation, thereby optimizing the available sequence space for coding regions, while relieving the circRNA-related size issue. This advancement broadens the repertoire of ORFs amenable to translation from circRNAs, enhancing their applicability as therapeutic agents in therapy.

## Methods

### Plasmids Construction

P-circ-3xF, described in Legnini *et al*. work^5^, was used as template for the new clones generated in this paper.

P-circ-3xF backbone vector was linearized through inverse PCR (Supplementary Table 1, primers 1-2); both GFP and EMCV amplification was performed on plasmid pcDNA3.1(+) eGFP (Addgene # 129020, Supplementary Table 1, primers 3-4, primers 5-6, respectively). In-Fusion Cloning (Takara) reaction was employed to generate the final EMCV-cGFP. From this vector, we removed EMCV IRES (Supplementary Table 1 primers 7 and 2) and replaced it with VCIP (amplified from genomic DNA, Supplementary Table 1, primers 8-9) or CVB3 IRES (amplified from Addgene plasmid #158645, Supplementary Table 1, primers 10-11) generating cGFP-VCIP and cGFP-CVB3, respectively.

In order to generate the TEE vectors, namely 13-glo-, 6.512- and 6.877-cGFP these elements, provided with terminal 15nt long sequences complementary to the backbone vector, were synthetized and annealed (Supplementary Table 1, oligos 12-13, oligos 16-17, oligos 18-19, respectively), and then inserted through In-Fusion Cloning (Takara) in the cGFP-EMCV vector linearized and devoid of the EMVC IRES (Supplementary Table 1, primers 14-15).

6.512-VCIP and 6.877-VCIP vectors were produced by linearizing the VCIP-cGFP vector (Supplementary Table 1, primers 21-22; Supplementary Table 1, primers 23-24, respectively), and amplifying 6.512 (Supplementary Table 1, primer 25-26) and 6.877 (Supplementary Table 1, primer 27-28) TEEs from the aforementioned synthetized corresponding elements to clone them through In-Fusion Cloning (Takara). eIF4G2 and PABP binding sequences combined with the 13mer positioned upstream (Supplementary Table 1, oligos 29-30 and oligos 31-32, respectively) were synthetized and annealed. cGFP-13-eIF4G and cGFP-13-PABP vectors were generated by inserting the above sequences in the cGFP-13-glo vector linearized devoid of the UTR using primers (table1 primer 1-2) through In-Fusion Cloning (Takara).

cGFP-UTR1, cGFP-UTR2 and cGFP-UTR3 were generated by linearizing the cGFP-13-glo vector using phosphorylated primers carrying circZNF609 UTR fragments. The linearized vector was ligated to reconstitute the vector. Specifically, cGFP-UTR1 was generated with (Supplementary Table 1, primers 33-34), cGFP-UTR2 was generated with (Supplementary Table 1, primers 35-36) and cGFP-UTR3 with (Supplementary Table 1, primers 37-38).

CircZNF609 containing clones were generated by linearizing p-circ-3xF vector (Supplementary Table 1, primers 39-40) in order to remove circZNF609 UTR and inserting either VCIP IRES (Supplementary Table 1, primers 41-42), 13-glo (Supplementary Table 1, annealed oligos 43-44) or 13-UTR3 (Supplementary Table 1 annealed oligos 45-46) in order to obtain VCIP-cZNF609, 13-glo-cZNF609, 13-UTR3-cZNF609 respectively.

All amplification reactions were carried out using CloneAmp mastermix polymerase (Takara). Ligase reaction was carried out using T4 DNA ligase enzyme, the reaction was kept at 16° for at least 15’ or more (up to 1h) based on the efficiency reached.

### Protein Analyses

Cells were harvested with 50-150 μL of Protein Extraction Buffer (100 mM Tris pH7.5, EDTA 1mM, SDS 2%, PIC1X (Complete, EDTA free, Roche) and incubated 10min on ice, then incubated on a rotator for 30min at 4°C and centrifuged at 13000xrpm for 10min at 4°C.

The supernatant was transferred to a clean tube, used or stored at −80°C. Total protein concentration was measured through the Bradford reagent (Bio-Rad Protein Assay) following manufacturer’s instructions.

25-50 μg of proteins were loaded on 4%–12% bis-tris-acrylamide gel (Life technologies) and transferred to a nitrocellulose membrane. The membrane was blocked in 5% milk and then hybridized with specific antibodies for 2hrs-o.n. at room temperature. After three washes in TBST, the filter was hybridized with the corresponding secondary antibody, if required, for one hour at room temperature. Protein detection was carried out with LiteAblot®EXTEND Long Lasting Chemiolominescent Substrate (EuroClone) or with Clarity Max Western ECL Blotting Substrate (BioRad) using ChemiDoc MP System and images were analyzed using Image Lab Software (BioRad). The antibodies used were Anti-Flag M2-Peroxidase (HRP) (Sigma-Aldrich, #A8592, 1:2500) and Anti-ACTB-Peroxidase (AC-15) monoclonal antibody (Sigma-Aldrich, #A3854, 1:2500).

### Cell culture

Human ERMS RD cell line (embryonal rhabdomyosarcoma cell line from a female patient) were cultured as in Dattilo *et al*.^11^. In particular, they were cultured in DMEM high glucose (Sigma-Aldrich) supplemented with 10% FBS (Sigma-Aldrich), l-glutammine (Sigma-Aldrich) 2 mM and penicillin-streptomycin 1× (Sigma-Aldrich).

SK-N-BE cells were cultured as described in Di Timoteo *et al*.^12^. In particular, cells were cultured in RPMI (Sigma-Aldrich, Saint Louis, MO, USA) supplemented with 10% FBS (Sigma-Aldrich, #F2442), GlutaMAX supplement 1X (Thermo Fisher Scientific, # 35050061), sodium-pyruvate 1 mM (Thermo Fisher Scientific, #11360070) and Pen/Strep 1X (Sigma-Aldrich, #P4458). All cells were cultured at 37 °C in a humidified atmosphere of 5% CO2.

For plasmids transfection, Cells (2,2x 10^5^) were plated in 2 ml medium in 35mm plates and transfected 24hrs later with plasmids (final amount 2,5µg) using 1,5 µL/mL of Lipofectamine 2000 Reagent (Thermo Fisher Scientific) and 75 µL/mL of Opti-MEM (Thermo Fisher Scientific). The medium was replaced 6-12hrs later. Cells were harvested 48hrs later.

For α-amanitin treatment, 7.5x 10^5^cells were plated in 60mm plates. After 24h each plate was split into two smaller plates, then either harvested (0h) or treated with α-amanitin (25 µg/ml) for 24h, before harvesting.

### RNA analysis

Total RNA in this study was extracted with Direct-zol RNA Miniprep (Zymo Research) kit according to the manufacturer’s specifications. DNase treatment was performed (Thermo Fisher Scientific).

Reverse transcription reactions of this study were performed with PrimeScript RT Master Mix (Takara Bio): 100-500ng of RNA were retrotranscribed in a 10 µL reaction mix with 1 µL of enzyme and 2 µL of buffer, incubated 10 min at 37°C and 5 min at 85C.

qPCR analyses in this study were performed with 4 ng equivalent of cDNA, 5 µL of 2X SYBR Mastermix (QIAGEN), 1 µL of 5 µM primers to a final volume of 10 µl. DNA amplification (40 cycles of 95C for 30 s, 55C for 30 s and 72C for 30 s, followed by melting curve analysis) was monitored with an ABI 7500 Fast or StepOnePlus System qPCR instrument

## Results

### A 63bp element combines efficiency and short size for circRNA translation

Several IRESs have been tested for their efficiency in circRNA translation^6^. Among them EMCV and CVB3, viral IRESs derived from the Encephalomyocarditis virus or Cocksakie virus respectively, have been described as two of the most proficient. Like viral ones, also cellular mRNAs can contain IRES elements^6,13^.

Nevertheless, cellular IRESs may differ from their viral counterparts in several features. Since secondary structure can impact on circularization, the fact that cellular IRESs appear to be less structured ^13^ makes them a good alternative to be applied to circRNAs.

We then firstly compared EMCV and CVB3 efficiency with the human IRES derived from vascular endothelial growth factor and type 1 collagen-inducible protein transcript (VCIP) in our system.

Such IRES has been reported to be ubiquitously active in human and mouse tissues and to be one of the most efficient among both human and viral IRESs to drive RNA translation^14 15^.

In order to test its efficiency in a circular context, we took advantage of a vector (cGFP) able to express a circularized flag-tagged GFP ORF (Fig.1A). The 3xFlag-tag, followed by a STOP codon, was inserted upstream the START codon so that it can be expressed only upon circularization (Fig. 1A). We cloned either EMCV, CVB3 or VCIP IRES between the initiation and the termination codons. In order to ensure to work with substantial RNA and protein yields, we tested these vectors in a tumor cell line, namely Rhabdomyosarcoma cells (RD). An empty vector was used as negative control (Ctrl).

**Fig. 1.**
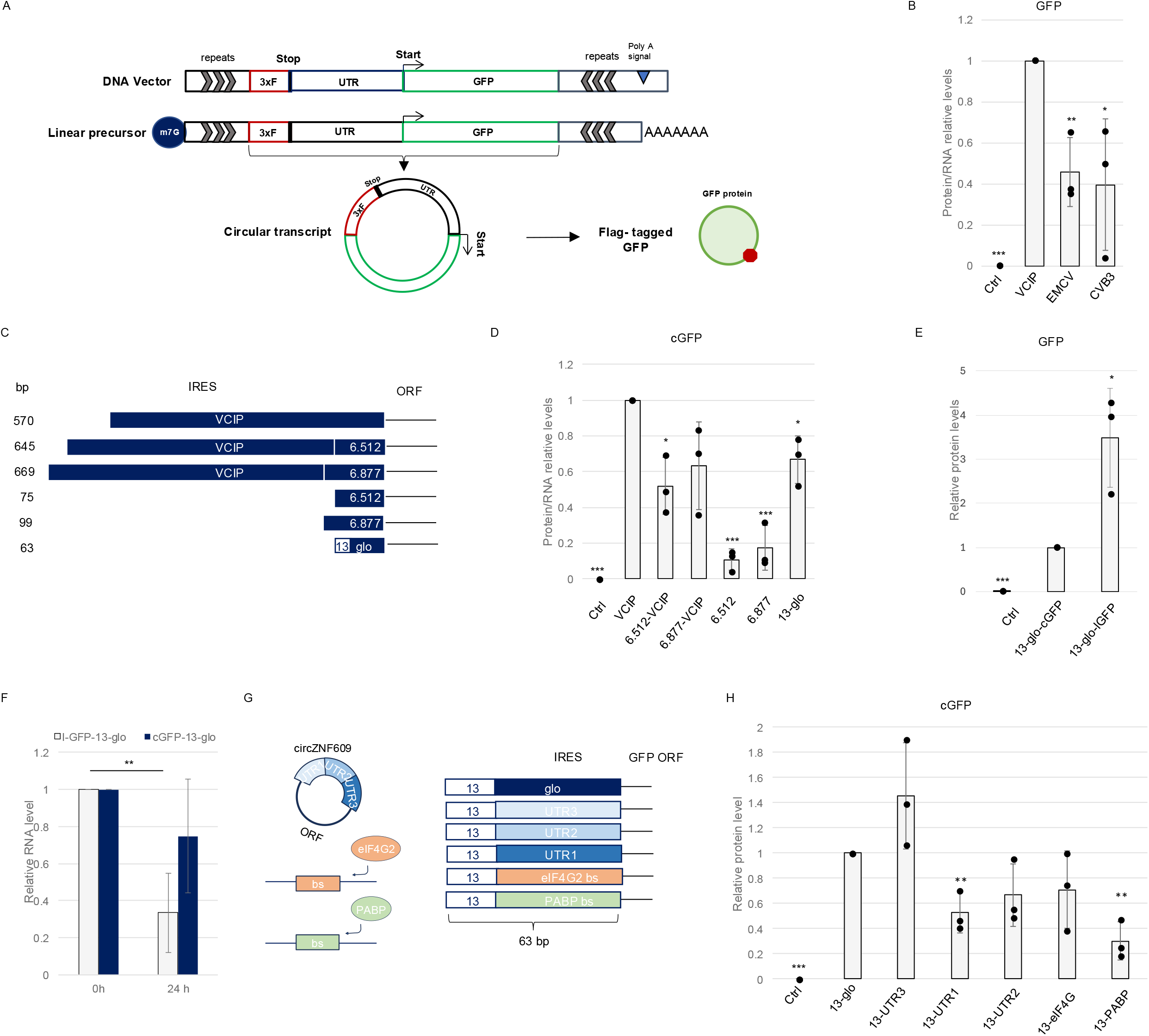
**A)** Schematic representation of the cGFP construct and the corresponding linear precursor RNA. The position of UTR (UTR), GFP ORF (GFP), Start/stop-codons, 3x-Flag-tag (3xF), inverted repeats flanking the circularization cassette (repeats), poly-A tail are indicated. **B)** Relative protein levels deriving from c-GFP translation and its IRESs mutant derivatives (VCIP, EMCV and CVB3) normalized on the corresponding transcript level. Levels were normalized over ACTB and expressed as relative quantities with respect to VCIP construct set to 1; Western blot are shown in fig. s1b. The ratio of each sample versus its experimental control was tested by two-tailed Student’s t test. * indicates a Student’s t test-derived p-value < 0.05, ** indicate a p-value < 0.01, and *** a p-value < 0.001. N=3. **C)** Schematic representation of VCIP, TEE and their derivatives. Length (bp) are indicated. **D)** Relative protein levels deriving from VCIP-cGFP translation and the indicated cGFP mutant derivatives normalized on the corresponding transcript level. Levels were normalized over ACTB and expressed as relative quantities with respect to VCIP construct set to 1; Western blot are shown in fig. s1g. The ratio of each sample versus its experimental control was tested by two-tailed Student’s t test. * indicates a Student’s t test-derived p-value < 0.05, ** indicate a p-value < 0.01, and *** a p-value < 0.001. N=3. **E)** Relative protein levels deriving from 13-glo-cGFP and 13-glo-lGFP translation. Levels were normalized over ACTB and expressed as relative quantities with respect to 13-glo-cGFP construct set to 1. The ratio of each sample versus its experimental control was tested by two-tailed Student’s t test. * indicates a Student’s t test-derived p-value < 0.05, ** indicate a p-value < 0.01, and *** a p-value < 0.001. N=3. **F)** Relative RNA levels of 13-glo-cGFP and its linear counterpart (13-glo-lGFP) at time point 0h (0H) or after 24h (24H) after alpha-amanitin treatment. Values are expressed as relative quantity with respect to time 0h set to a value of 1. N=3. **G)** Schematic representation of 13-glo and its spacer variants: β-glo UTR fragment originally used as spacer has been replaced with either single circZNF609 UTR fragments (blue) or eIF4G2 (orange)/PABP (green) binding site.. Length (bp) are indicated. **H)** Relative protein levels deriving from 13-glo-cGFP and its indicated spacer mutants derivativas normalized on the corresponding cGFP transcript level. Levels were normalized over ACTB and expressed as relative quantities with respect to 13-glo-cGFP construct set to 1. The ratio of each sample versus its experimental control was tested by two-tailed Student’s t test. * indicates a Student’s t test-derived p-value < 0.05, ** indicate a p-value < 0.01, and *** a p-value < 0.001. N=3.

The EMCV and the VCIP vectors produced comparable amount of mature circRNA (Fig. S1A), with VCIP producing a 2-fold increase in protein production (fig. S1B). CVB3, instead, possibly due its extensive length and structure^16^ negatively impacted on circular GFP transcript level making a direct protein level comparison difficult (fig. S1A). When normalizing the derived protein on circRNA level, VCIP resulted the most proficient IRES in RD cells (fig. 1B).

We also checked for precursor RNA levels produced from the vectors under analyses. While producing comparable amounts of circular transcripts, more precursor RNA derived from VCIP when compared to EMCV (fig. S1C). Instead, CVB3, showed a drastic drop paralleling the drop in circRNA level (fig. S1C). These data suggest that the different IRESs impact on the precursor transcript levels produced from these vectors. In conclusion we demonstrated that VCIP IRES is more efficient with respect to viral IRESs such as EMCV and CVB3 in driving the translation of a GFP circular mRNA.

However, the employment of IRESs extended sequences for translation does not appear to be an optimal choice when considering circRNAs as therapeutic tools. Indeed, IRESs length has been reported to significantly increase secondary structure formation^17^. In addition, long circRNAs are more prone to nicking^18^.

Thus, we employed Translation Enhancing Elements (TEEs), small size sequences that can be found at the 5’ end of a transcript, to test their ability to drive circRNA translation. TEEs have been demonstrated to enhance mRNAs translation, and their activity has been studied in linear transcripts only, yet^10^.

In particular, we selected different TEEs based on the efficiency reported and we adopted them as cGFP UTRs (fig. 1C). Since TEEs activity relies on the presence of upstream START codons, we ensured that no significant ORFs (>20 aminoacids) were generated by TEEs insertion upstream the GFP ORF.

A very brief sequence (13nt, “13-mer”) has been described as a common feature among TEEs. Such 13-mer element works at its best if properly spaced (about 50 nt) from the ORF START codon in a linear context^10^. Aiming to combine efficiency and small size, we then adopted this element as UTR in the cGFP vector, spaced from the START codon by a randomly selected fragment from β-Globin UTR (Fig. 1C).

Furthermore, we also added TEEs elements upstream the VCIP IRES in the vector VCIP-cGFP in order to check any synergistic activity of the two elements (VCIP and TEEs) and eventual substantial advantages in translation even at the expense of scale advantages.

As shown in fig. S1D, the analysis of the mature circRNA revealed that we obtained different levels of circular transcript from different vectors. However, we did not detect strong differences in circGFP levels deriving from VCIP-cGFP and its TEE mutant derivatives (VCIP-6.512 and VCIP-6.877).

Instead, a 2-fold increase in the RNA amount was observed in the vectors devoid of VCIP with respect to their IRES-containing counterparts (fig. S1d), possibly due to the smaller size of the circular RNA produced. Indeed, about 500-bp of the VCIP IRES are absent from the cGFP produced by 6.877, 6.512 or 13-glo (fig. 1C). We investigated whether differences in circRNAs levels derived from the vectors under study might be due to differences in precursors RNA levels.

Despite the circular transcript increase (fig. S1D), no significant increase in precursor RNA level was detected with the exception of 6.512-VCIP candidate (fig. S1E). Moreover, all the new candidate vectors apart from 13-glo showed a decreased translation ability in comparison to the VCIP-cGFP used as reference (fig. S1F, fig. S1G), excluding them as circRNA translation enhancers. Notably, when normalizing the amount of protein produced and the respective RNA level, 13-glo stood out as an element possibly combining the ability to efficiently drive circRNA translation with its very short size (fig. 1D, fig. S1F, fig. S1G).

To better compare the translation efficiency of VCIP and 13-glo, we then tried to make the level of cGFP transcript comparable by adjusting the amount of plasmid transfected in the cells (Fig. S1H). Importantly, upon such adjustments, we found that cGFP protein signal from 13-glo-cGFP was comparable to VCIP-cGFP, but with considerable advantages in terms of size (fig. S1I).

These results led us to elect 13-glo as our best candidate, since combining efficiency and solving size issues, beside ensuring higher circRNA level.

However, as expected, when comparing the cap-dependent translation and the 13-glo-driven cap-independent translation of the same GFP ORF coded by a linear or circular RNA respectively, cap-independent translation resulted as less efficient (fig. 1E).

Indeed, despite the fact that linear RNA was expressed more than 10 folds less if compared to the circular one (fig. S1J), it could produce about 3-folds more protein (fig. 1E).

If it is true that circRNA translation resulted still much more inefficient than translation from a linear mRNA, it has also been demonstrated that circRNA, being more resistant to exonucleases activity, last longer than their linear counterparts ensuring protein production for a longer period even if at lower levels ^19,20^. We validated such results also for 13-glo-derived transcripts (fig. 1F). Coherently, when comparing the persistency of the transcripts derived from vectors producing either a circular (cGFP-13-glo) or linear GFP transcript (lGFP-13-glo) upon transcription inhibition obtained through α-amanitin treatment, we observed that after 24hrs treatment the linear transcript experienced a drastic drop (70%) when compared to the circular one, remaining stable (fig.1F).

In conclusion, we demonstrated that VCIP, a human IRES, and a 13-nt short sequence element (13-glo), are able to efficiently drive the translation from a circular transcript comparably to commonly adopted IRES such as EMCV and CVB3 (about 600-800 nt) providing a clear length advantage to be exploited during *in vitro* circRNA synthesis and expanding the repertoire of ORFs potentially suitable for being translated from a synthetic circRNA.

Since 13-mer works best if properly spaced from the START codon, we wondered whether the spacer choice could affect its activity. With this purpose we replaced the initially randomly adopted 50nt β-globin UTR fragment with sequences potentially able to tether on cGFP transcript factors involved in translation, namely sequences recruiting either eIF4G (eIF4G aptamer) or PABP (poly-A binding protein), previously reported to enhance IRES activity^21^.

Furthermore, looking for alternative elements, we opted for the UTR of circZNF609. Such UTR is about 150 nt long and an optimal spacer has been reported to be of 50 nt.

We arbitrarily fragmented circZNF609 UTR into three 50-nt fragments and used them as spacer in the vectors we named 13-UTR1, 13-UTR2, and 13-UTR3.

The insertion of such elements did not induce changes in the RNA level in any case but for PABP (13-PABP), which showed a trend of upregulation if compared to our best candidate 13-glo (fig. S1L).

However, none of the new candidates could produce a higher amount of protein with respect to 13-glo with the only exception of 13-UTR3, which showed an average 1.5-fold increase if compared to 13-glo (fig. 1G, fig. S1L), showing better performance even than VCIP-cGFP that show only a 1.2-fold increase with respect to the same reference.

In conclusion, we showed that spacer sequence can affect 13-mer translation activity and in particular UTR3 of circZNF609 can positively impact on protein production from circular mRNA.

These results indicate that possibly future high-throughput screenings might identify spacer able to further boost circRNA translation.

### 13-mer translation efficiency is ORF and cell-type independent

In order to check if the translation efficiency of the elements we tested could be dependent on the downstream ORF, we tested VCIP, the most efficient among the IRESs selected, as well as 13-glo and 13-UTR3, the most promising among the TEE-derived UTR, on the ORF of circZNF609, an endogenous translated circRNA^5^. With this purpose we produced the vectors VCIP-cZNF609, 13-glo-cZNF609, and 13-UTR3-cZNF609. Despite variations on circRNA levels produced by the vectors (fig. S2A), almost paralleled by variations in their precursor level (fig. S2B), all our candidates better drive translation if compared to the natural circZNF609 UTR (ZNF) (fig. 2A, fig. S2C). Among them, despite slight differences in RNA levels, TEE-derived UTRs were confirmed to be more efficient than VCIP-cZNF609, with 13-UTR3-cZNF609 being the most efficient in circRNA global translation (fig. 2A; fig. S2C).

**Fig. 2.**
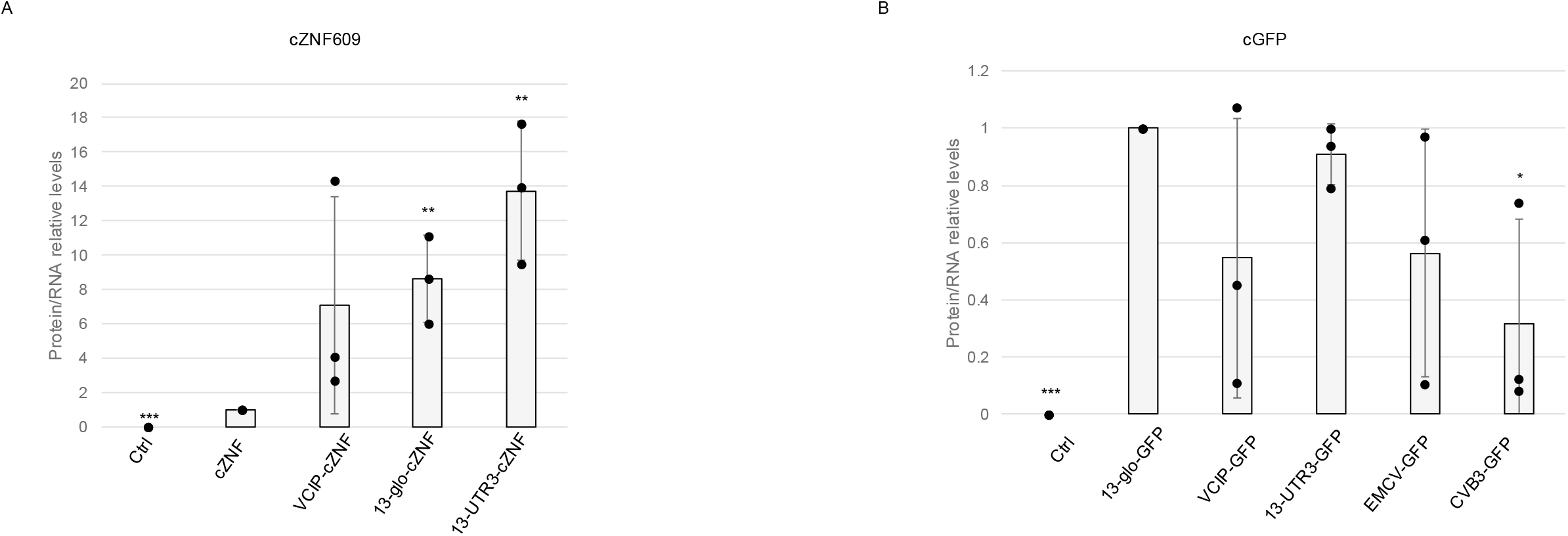
**A)** Relative protein levels deriving from c-ZNF609 and its indicated mutant derivatives normalized on the corresponding transcript level. Levels were normalized over GAPDH and expressed as relative quantities with respect to c-ZNF609 construct set to 1; Western blot are shown in fig. s2c. The ratio of each sample versus its experimental control was tested by two-tailed Student’s t test. * indicates a Student’s t test-derived p-value < 0.05, ** indicate a p-value < 0.01, and *** a p-value < 0.001. N=3. **B)** Relative protein levels deriving from c-GFP and its indicated mutant derivatives normalized on the corresponding transcript level in SK-N-BE cells. Levels were normalized over ACTB and expressed as relative quantities with respect to 13-glo-cGFP construct set to 1; Western blot are shown in fig. s2e. The ratio of each sample versus its experimental control was tested by two-tailed Student’s t test. * indicates a Student’s t test-derived p-value < 0.05, ** indicate a p-value < 0.01, and *** a p-value < 0.001. N=3.

These results suggest that the efficiency of the elements tested is not ORF-dependent but also that such elements show far better performances than natural circZNF609 UTRs.

Since circZNF609 is translated into two proteins (30 kDa and 35 kDa) from two alternative START codons, we also analyzed the translation levels of each protein product individually. As shown in Fig. S2D, both TEE derivatives (13-glo and 13-UTR3) drive translation from the first START codon (producing the 35 kDa protein) more efficiently than VCIP. However, when assessing translation from the second ATG (producing the 30 kDa protein), 13-glo is less efficient than both VCIP and 13-UTR3, though still markedly more efficient than the natural circZNF609 UTRs. This result becomes particularly relevant in the uncommon scenario where the expression of a protein initiated from two alternative start codons is desired. In such a context, the data presented here indicate that 13-UTR3 represents the most suitable element to ensure efficient translation from both initiation sites.

Another aspect worth testing was whether the cellular context in which we had tried our candidates could impact on their efficiency, so we changed cellular model and tested the ability of our candidates in driving cGFP translation in SK-N-BE cells, a neuroblastoma cell line. We tested both the IRESs we previously showed (VCIP, EMCV and CVB3) and our best short candidates (13-glo and 13-UTR3). As shown in fig. S2E, we confirmed the trend in RNA level previously observed, in fact 13-glo resulted in a 2-fold increase and 13-UTR3 a 3-fold increase when compared to VCIP (fig. S2E). EMCV and CVB3, instead, showed a significant decrease in circRNA level. Coherently with our previous results, in terms of protein production, 13-glo and 13-UTR3 were the most efficient compared to the other elements tested also in SK-N-BE (fig. 2B, fig. S2F).

## Discussion

The potential of RNA-based drugs was widely demonstrated during the COVID-19 pandemic. In addition to vaccines, many other diseases can be tackled through the use of RNA technologies. This is supported by the large number of preclinical and clinical studies employing them. A prominent example is cancer vaccines, but accumulating evidence suggests that RNA-based drugs can be successfully used in other diseases, such as cystic fibrosis, autoimmune diseases, metabolic diseases, cardiovascular diseases^22–27^.

Despite this, the use of linear RNA molecules for therapeutic purposes is strongly hindered by their instability, high immunogenicity, high storage costs, and difficulties in large-scale production, also due to the need of chemical modification addition needed for their stabilization ^17^.

CircRNAs do not suffer from many of the typical limitations of linear RNAs and have thus attracted attention in the field of RNA therapies. Indeed, due to their intrinsic structure, circRNAs exhibit high stability and RNase resistance^19^, conferring them a significantly longer half-life than their linear RNA counterparts^28^ and allowing their storage at up to 37 °C for up to 28 days^29^. Moreover, the coding longevity of artificially translatable circRNAs was demonstrated to be much greater than that of their nucleoside-modified linear mRNA equivalents^30–33^. All these advantages are particularly important when considering the adoption of RNA-based therapies.

Given these considerations, circRNAs may potentially represent the next generation of RNA-based drugs.

CircRNAs have been already proposed as vaccines against COVID-19 following preclinical studies demonstrating their effectiveness in mice and macaques^34^. Indeed, a circRNA-based vaccine was constructed to encode SARS-CoV-2-specific neutralizing antibodies to achieve COVID-19 prevention. Furthermore, an *in vivo* study of circRNA-based vaccines for the treatment and prevention of tumors (melanoma) has also been conducted, showing its strong prophylactic and anti-metastatic potential^33–37^. Moreover, *in vivo* NGF expression from a circular RNA was effective in providing neuroprotection in mouse retinal ganglion cells under glucose starvation ^20^. However, further *in vivo* and clinical studies will be essential for their effective development and therapeutic application, as well as there is the urgent need to relief circRNAs weaknesses.

Many of the limitations associated with circRNA use are mainly related to their length and their ability to be translated, which impact their production and efficiency, respectively^38–40^. For this reason, there is a common belief in the field that the first step in engineering circRNAs should aim to resolve or mitigate these specific issues, in order to find a compromise for a circRNA to encode an average ORF, but also to maintain a suitable size.

Shortening circRNAs is considered a focal point for their adoption as therapeutic tools. In the field, considerable effort has been directed towards reducing the ORFs, given the inability to shorten the length of known IRESs without compromising their efficiency. This is why many studies aimed to produce short peptides from circRNAs^41^.

However, this strategy narrows down the choice of proteins suitable for circRNA-mediated translation and, consequently, the range of possible coding circRNAs-based therapeutic approaches. To date, the most commonly elements used to drive circRNA translation are internal ribosome entry sites (IRES), particularly viral ones. However, these sequences tend to be very long (600-1000 nt) and structured, and their efficiency depends on cell type and species. Currently, CVB3, a viral IRES, is the most used for circRNA translation^34 20^. In this study, we compared it with other IRESs both in a muscle-(RD) and in neural-like cellular model (SK-N-BE). Importantly, one of these, VCIP, is a human IRES, ubiquitously expressed, and was found to be more efficient in promoting circRNA translation in a cell type- and ORF-independent manner ^15^.

Nevertheless, VCIP is approximately 600 nt long. Thus, as other IRESs, even ensuring efficient RNA translation, it is not solving the size issue. Given the importance of shortening of the elements that allow circRNA translation we focused on the identification of shorter sequences.

In this study we identified a very short element (approximately 60nt) able to drive circRNA translation more efficiently than commonly adopted IRESs (600-800 nt) with a considerable advantage in terms of size. This element consists of a 13-nt sequence previously identified as common feature in linear mRNAs TEEs ^10^ and a spacer. We explored the possibility to further enhance its performance by varying the sequence of the spacer through a rational approach considering elements known to recruit translation factors and sequences derived from the UTR of an endogenously translated circRNA (circZNF609)^5^. Our results indicated that a fragment of the UTR of circZNF609 (UTR3) is able to improve the translation. Nevertheless, high-throughput approaches, not in the scope of this manuscript, may be used in the future in order to find even more proficient elements and further optimize circRNA translation driven by 13-mer.

Also, circRNA translation driven by 13-mer might be ameliorated by combining its power with other approaches not affecting the circRNAs size, such rolling translation, covalent or non-covalent attachment of an m^7^G cap or by exploring the effects of RNA modifications (e.g. m6A or *N*^1^-methylpseudouridine) with the aim to increase circRNA potential as therapeutic tools.

Our data suggest that the efficiency of 13mer-UTR3 is independent from the ORFs and from the cell type where the circRNA is expressed. These data do not exclude the possibility that further study will identify elements driving circRNA translation in a tissue-specific manner in order to enhance the specificity of potential circRNA-based drugs.

In conclusion, we massively reduced the length of regulatory elements driving circRNA translation, providing the possibility to save space for coding sequences, expanding the landscape of ORFs potentially suitable for being translated from circRNAs and serve as therapeutic circular RNAs in the treatment of diseases.

## Supporting information

Suppemental Table 1

## Acknowledgements

We thank F. Margarita and M. Marchioni for technical help and M. Caruso for assistance. PcDNA CVB3 was a gift from Kevin Janes (Addgene plasmid # 158645). PcDNA3.1(+) eGFP was a gift from Jeremy Wilusz. This work was partially supported by grants from ERC-2019-SyG 855923-ASTRA, AIRC IG 2019 Id. 23053, and PRIN 2017 2017P352Z4 to I.B.; “National Center for Gene Therapy and Drug based on RNA Technology” (CN00000041) and NextGenerationEU PNRR MUR to I.B.; Sapienza departmental projects 2023 (RD12318A998C70B8) to G.D.T.

---

**Fig. S1.**
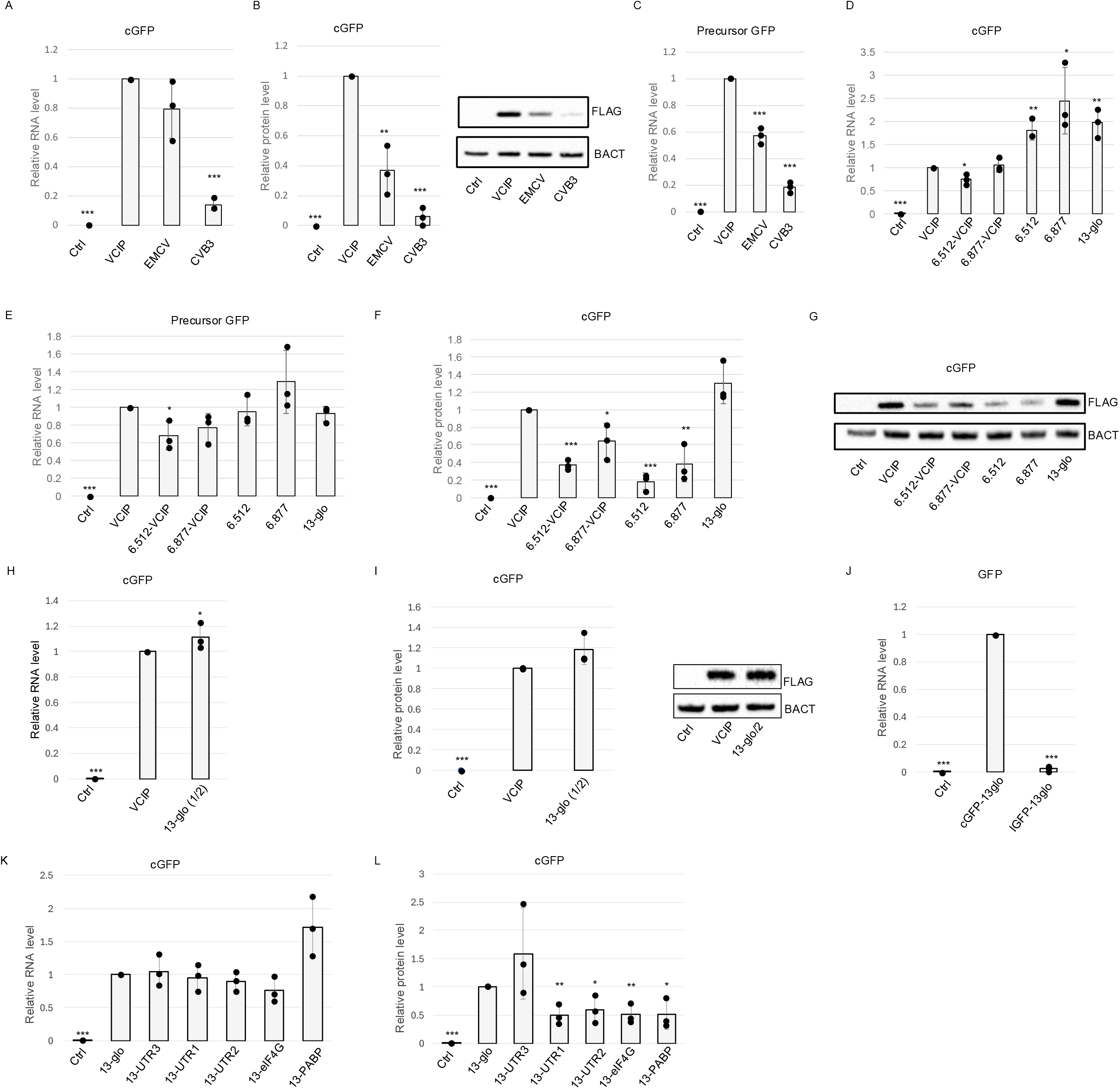
**A)** Relative RNA levels of circular transcripts from the indicated cGFP IRES mutant derivatives are shown. Values were normalized against GAPDH and expressed as relative quantity with respect to VCIP-cGFP set to a value of 1. The ratio of each sample versus its experimental control was tested by two-tailed Student’s t test. * indicates a Student’s t test-derived p-value < 0.05, ** indicate a p-value < 0.01, and *** a p-value < 0.001. N=3. **B)** Relative protein levels deriving from c-GFP translation and its IRESs mutant derivatives (VCIP, EMCV and CVB3). Levels were normalized over ACTB and expressed as relative quantities with respect to VCIP construct set to 1 (left panel). The ratio of each sample versus its experimental control was tested by two-tailed Student’s t test. * indicates a Student’s t test-derived p-value < 0.05, ** indicate a p-value < 0.01, and *** a p-value Student’s t test-derived p-value < 0.05, ** indicate a p-value < 0.01, and *** a p-value < 0.001. N=3. < 0.001. N=3. A representative Western blot is shown (right panel). **C)** Relative RNA levels of precursor transcripts from the indicated cGFP IRES mutant derivatives are shown. Values were normalized against GAPDH and expressed as relative quantity with respect to VCIP-cGFP set to a value of 1. The ratio of each sample versus its experimental control was tested by two-tailed Student’s t test. * indicates a Student’s t test-derived p-value < 0.05, ** indicate a p-value < 0.01, and *** a p-value < 0.001. N=3. **D) E)** Relative RNA levels of circular (D) and precursor (E) transcripts from VCIP-cGFP and the indicated cGFP TEE mutant derivatives. Values were normalized against GAPDH and expressed as relative quantity with respect to VCIP-cGFP set to a value of 1. The ratio of each sample versus its experimental control was tested by two-tailed Student’s t test. * indicates a Student’s t test-derived p-value < 0.05, ** indicate a p-value < 0.01, and *** a p-value < 0.001. N=3. **F) G)** Relative protein levels deriving from VCIP-cGFP translation and its indicated TEEs mutant derivatives (F). Levels were normalized over ACTB and expressed as relative quantities with respect to VCIP construct set to 1 (left panel). The ratio of each sample versus its experimental control was tested by two-tailed Student’s t test. * indicates a Student’s t test-derived p-value < 0.05, ** indicate a p-value < 0.01, and *** a p-value < 0.001. N=3. A representative Western blot is shown (G). **H) I)** Relative RNA (H) and protein (i, left) levels detected upon trasfection of VCIP-cGFP vector and half of the amount of 13-glo-cGFP. Values were normalized against GAPDH or ACTNB for RNA and proteins, respectively, and expressed as relative quantity with respect to VCIP-cGFP set to a value of 1. The ratio of each sample versus its experimental control was tested by two-tailed Student’s t test. * indicates a Student’s t test-derived p-value < 0.05, ** indicate a p-value < 0.01, and *** a p-value < 0.001. N=3. A representative Western blot is shown (i, right panel). **J)** Relative RNA levels deriving from 13-glo-cGFP and 13-glo-lGFP. Levels were normalized over GAPDH and expressed as relative quantities with respect to 13-glo-cGFP construct set to 1. The ratio of each sample versus its experimental control was tested by two-tailed Student’s t test. * indicates a Student’s t test-derived p-value < 0.05, ** indicate a p-value < 0.01, and *** a p-value < 0.001. N=3. **K) L)** Relative RNA (K) and protein (L) levels deriving from 13-glo-cGFP and its indicated spacer mutants derivativas. Levels were normalized over GAPDH or ACTNB for RNA and protein, respectively,and expressed as relative quantities with respect to 13-glo-cGFP construct set to 1. The ratio of each sample versus its experimental control was tested by two-tailed Student’s t test. * indicates a Student’s t test-derived p-value < 0.05, ** indicate a p-value < 0.01, and *** a p-value < 0.001. N=3.

**Fig. S1.**
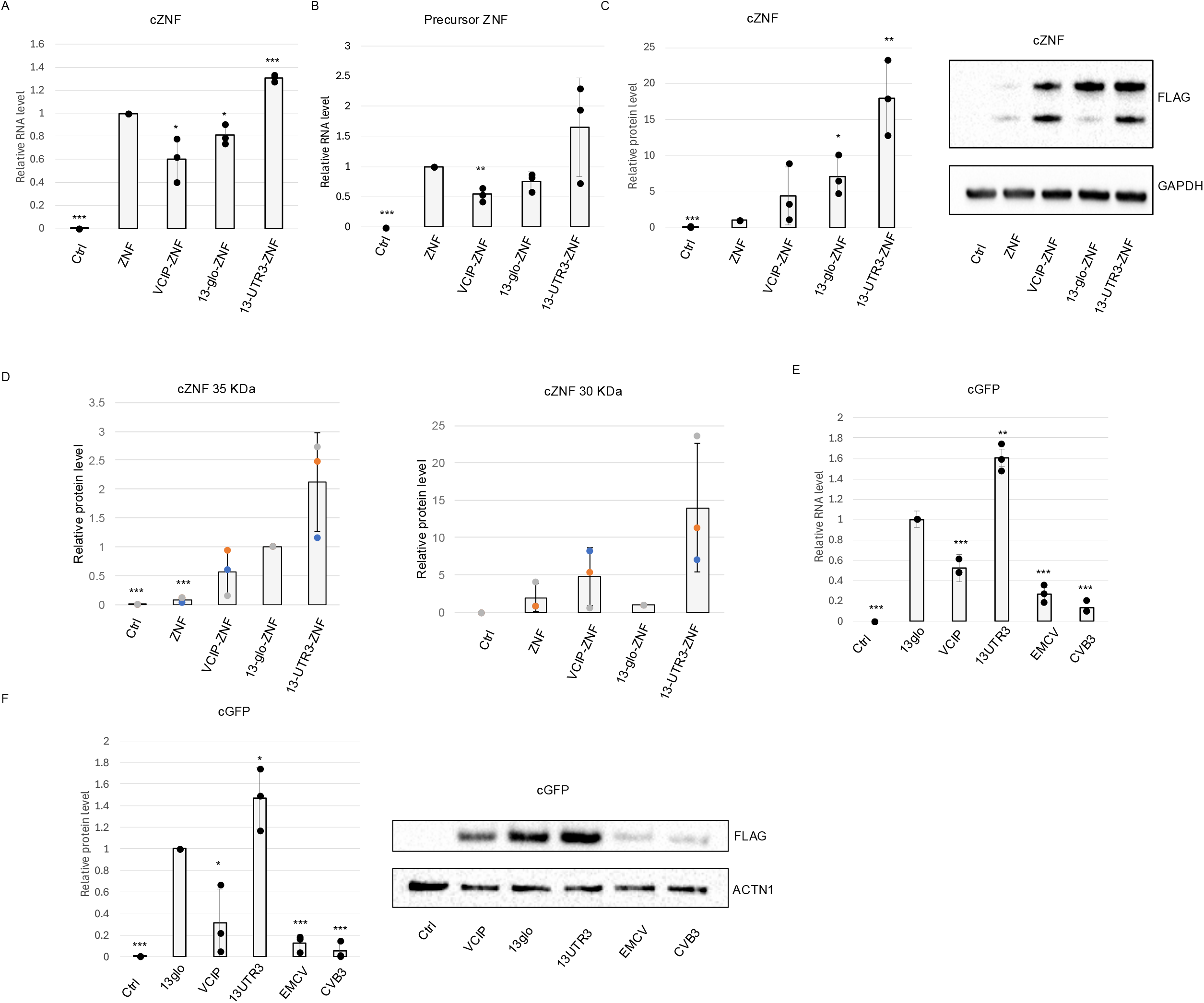
**A) B)** Relative RNA levels of either circular transcripts (A) or its precursor (B) poroduced from cZNF609 and the indicated mutant derivatives. Values were normalized against GAPDH and expressed as relative quantity with respect to circZNF609 set to a value of 1. The ratio of each sample versus its experimental control was tested by two-tailed Student’s t test. * indicates a Student’s t test-derived p-value < 0.05, ** indicate a p-value < 0.01, and *** a p-value < 0.001. N=3. **C)** Relative protein levels deriving from circZNF609 translation and its indicated mutant derivatives. Levels were normalized over GAPDH and expressed as relative quantities with respect to cZNF construct set to 1 (left panel). The ratio of each sample versus its experimental control was tested by two-tailed Student’s t test. * indicates a Student’s t test-derived p-value < 0.05, ** indicate a p-value < 0.01, and *** a p-value < 0.001. N=3. A representative Western blot is shown (right panel). **D)** Relative protein levels deriving from each circZNF609 ORF. Levels were normalized over GAPDH and expressed as relative quantities with respect to 13-glo-cZNF construct set to 1. Replicates are identified by color. **E) F)** Relative RNA (D) ad protein (e, left panel) levels deriving from cGFP and its indicated mutant derivatives in SK-N-BE cells. RNA or proteins kevels were normalized over GAPDH or ACTN1, respectively, and expressed as relative quantities with respect to 13-glo-cGFP construct set to 1; The ratio of each sample versus its experimental control was tested by two-tailed Student’s t test. * indicates a Student’s t test-derived p-value < 0.05, ** indicate a p-value < 0.01, and *** a p-value < 0.001. N=3. A representative Western blot is shown (e, right panel)

