## Supplementary material for "A Short 63-Nucleotide Element Promotes Efficient circRNA Translation": Suppemental Table 1

| Oligonucleotides and other sequence-based reagents | Sequence |
|---|---|
| 1fw | TCACTTGTCATCGTCATCCT |
| 2 rev | GTAAGAAGCAAGGTTTCATTTAGGG |
| 3EGFP FW | AATATGGCCACAaccatggtgagcaagggcgaggag |
| 4EGFP REV | AACCTTGCTTCTTACcttgtacagctcgtccatgc |
| 5EMCV FW | GACGATGACAAGTGAatccGCCCCTCTCCCTCCCC |
| 6EMCV REV | ggtTGTGGCCATATTATCATCGTGTTTTTCAAAGGAAAAC |
| 7Inv p-circ ires fw | atggtgagcaagggcgaggag |
| 8VCIP infusion fw | AGGATGACGATGACAAGTGAgACCTCGTGAAATAAAAGTGC |
| 9VCIP inf rev | tcctcgcccttgctcaccatATGGCGCTGGCTGCGGCGCG |
| 10Cvb3 inf fw | GACGATGACAAGTGAgagacccaagcttggtacc |
| 11Cvb3 inf rev | gcccttgctcaccattttgctgtattcaacttaac |
| 12Beta glo FW | GACGATGACAAGTGAAAATCAATAAATGACATTTGCTTCTGAC ACAACTGTGTTCACTAGCAACCTCAAACAGACACC |
| 13Beta glo REV | gcccttgctcaccatGGTGTCTGTTTGAGGTTGCTAGTGAACACAGT TGTGTCAGAAGCAAATGTCATTTATTGATTT |
| 14Inv glo fw | atggtgagcaagggcgagg |
| 15Inv glo rev | TCACTTGTCATCGTCATCCT |
| 16 Fw 512 | AGAGAAATGCAAATCAAAACCACAATGGAATACCATCTCACG CCAGTCAGAATGGCAATTATTAAAAAATCACAACAATTAATGA TGGCAAGGCTGTGG |
| 17 Rev 512 | gcccttgctcaccatCCACAGCCTTGCCATCATTAATTGTTGTGATTT TTTAATAATTGCCATTCTGACTGGCGTGAGATGGTATTCCATT GTGGTTTTGATTTGCATTTCTCTTCACTTGTCATCGTC |
| 18 Fw 877 | GACGATGACAAGTGAAAAGAAATGGAATCGAAGAGAATGGAA ACAAATGGAATGGAATTGAATGGAATGGAATTGAATGGAATG GGAACGatggtgagcaagggc |
| 19 Rev 877 | gcccttgctcaccatCGTTCCCATTCCATTCAATTCCATTCCATTCAA TTCCATTCCATTTGTTTCCATTCTCTTCGATTCCATTTCTTTTC ACTTGTCATCGTC |
| 20 VCIP- 6.512 FW | GACCTCGTGAAATAAAAGTGCAG |
| 21 VCIP- 512 FW | ATGGCAAGGCTGTGGGACCTCGTGAAATAAAAGTG |
| 22 VCIP-512 REV | GATTTGCATTTCTCTTCACTTGTCATCGTCATCCTTGTA |
| 23 VCIP- 6.877 FW | AATGGAATGGGAACGGACCTCGTGAAATAAAAGTG |
| 24 VCIP- 6.877 REV | CGATTCCATTTCTTTTCACTTGTCATCGTCATCCT |
| 25 512 FW | AGAGAAATGCAAATCAAAACC |
| 26 512 REV | CCACAGCCTTGCCATCAT |
| 27 877 FW | AAAGAAATGGAATCGAAGAGAATG |
| 28 877 REV | CGTTCCCATTCCATTCAATTC |
| 29 IF4G FW | GACGATGACAAGTGAAAATCAATAAATGACTCACTATTTGTTT TCGCGCCCAGTTGCAAAAAGTGTCGTAATTGACTAAatggtgagc aagggc |

| | |
|---|---|
| 30 IF4GREV | gcccttgctcaccatTTAGTCAATTACGACACTTTTTGCAACTGGGCGCGAAAACAAATAGTGAGTCATTTATTGATTTTCACTTGTCATCGTC |
| 31 PABP FW | GACGATGACAAGTGAAAATCAATAAATGAAAAAAAAAAAACCAAAAAAAAAAAACAAAAAAAAAAAAATAATTGACTAAatggtgagcaagggc |
| 32 PABP REV | gcccttgctcaccatTTAGTCAATTATTTTTTTTTTTTGTTTTTTTTTTTTGGTTTTTTTTTTTTCATTTATTGATTTTCACTTGTCATCGTC |
| 33 UTR1 FW | CAAGGAGGCAGGTCTGAGAACGTAGatggtgagcaagggcgaggagctgt |
| 34 UTR1 REV | GTCAGAACATGTCCAGTGGACAACACATTTATTGATTTTCACTTGTCATC |
| 35 UTR2 FW | AAAATAGGAAAGCTGGGGGCAAGGAatggtgagcaagggcgaggagctgt |
| 36 UTR2 REV | CAGCTTCAGCTACGTTCTCAGACCTCATTTATTGATTTTCACTTGTCATC |
| 37 UTR3 FW | TTGAACCTTGAGGTGGGACGTTGACTCTAAGatggtgagcaagggcgagg |
| 38 UTR3 REV | GGCTCTTCCTTGCCCCCAGCATTTATTGATTTTCACTTGTCATCGTCATCC |
| 39 INV ZNF FW | ATGTCCTTGAGCAGTGGAGC |
| 40 INV ZNF REV | TCACTTGTCATCGTCATCCTTG |
| 41 ZNF VCIP FW | GACGATGACAAGTGAGACCTCGTGAAATAAAAGTG |
| 42 ZNF VCIP REV | ACTGCTCAAGGACATATGGCGCTGGCTGCGGCGCG |
| 43 ZNF 13BETA FW | GACGATGACAAGTGAAAATCAATAAATGACATTTGCTTCTGACACAACTGTGTTCACTAGCAACCTCAAACAGACACCATGTCCTTGAGCAGT |
| 44 ZNF 13BETA REV | ACTGCTCAAGGACATGGTGTCTGTTTGAGGTTGCTAGTGAACACAGTTGTGTCAGAAGCAAATGTCATTTATTGATTTTCACTTGTCATCGTC |
| 45 ZNF 13UTR3 FW | GACGATGACAAGTGAAAATCAATAAATGCTGGGGGCAAGGAAGAGCCTTGAACCTTGAGGTGGGACGTTGACTCTAAGATGTCCTTGAGCAGT |
| 46 ZNF 13UTR3 REV | ACTGCTCAAGGACATCTTAGAGTCAACGTCCCACCTCAAGGTTCAAGGCTCTTCCTTGCCCCCAGCATTTATTGATTTTCACTTGTCATCGTC |
| qPCR primers | |
| 47 gfp cntr fw | TGAGCAAAGACCCCAACGAG |
| 48 flag rev | GTCATCGTCATCCTTGTAATCGA |
| 49 hung rev | ACAGGCACTCAGCAGCACAAA |
| 50 gapdh fw | CCAAAATCAAGTGGGGCGAT |
| 51 gapdh rev | GGCAGAGATGATGACCCTTT |
| 52 Znf real fw | GAGGAAGGGGAGAATGAGTG |
